# Genome-Wide Association Study Identifies Novel Genetic Variants Linked to Meropenem Resistance in *Klebsiella pneumoniae*

**DOI:** 10.64898/2026.11.24.768875

**Authors:** Ojas Bardiya, Anne Clatworthy, Josephine Bagnall, Christiaan Wijers, Vincenzo di Natale, Jack Shigeta, Noam Shoresh, Deborah Hung

## Abstract

Carbapenem-resistant Enterobacteriaceae have emerged as an urgent public health threat in recent years. Infections caused by carbapenem-resistant *Klebsiella pneumoniae* in particular are associated with high rates of mortality, especially in immunocompromised patients. Substantial carbapenem resistance remains unexplained in *K. pneumoniae* even after accounting for known resistance genes. To uncover unknown mechanisms associated with meropenem resistance, we performed a genome-wide association study (GWAS) in 1,705 *K. pneumoniae* clinical isolates with previously determined resistance profiles to meropenem . We conducted the GWAS by using an elastic net regression to control for genome-wide linkage disequilibrium and combined it with a conditional linear mixed model to control for the presence of genes known to confer carbapenem resistance. The resulting set of variants predicted to be associated with meropenem resistance mapped to genes broadly related to the plasticity of the genome. We developed a visualization tool integrating GWAS results with phenotypic data and polymorphic site information to aid in hypothesis generation, which allowed us to prioritize novel genomic candidates. Our results demonstrate the effectiveness of the combined approach in detecting novel genomic variants associated with meropenem resistance in *K. pneumoniae*, while controlling for strong bacterial population structure, to advance our understanding of the genetic basis of carbapenem resistance.

## INTRODUCTION

The rise of carbapenem resistance in Enterobacteriaceae has been recognized as a critical public health threat by the World Health Organization^1,2^. Carbapenem-resistant *Klebsiella pneumoniae* (CRKp), in particular, is a major source of multi-drug resistant nosocomial infections and is associated with high mortality rates due to limited effective therapeutic options^11/24/2026 2:15:00 PM^. The increasing incidence of CRKp is partly attributed to the acquisition of *K. pneumoniae* carbapenemase (KPC) genes such as bla_KPC_, bla_NDM_, bla_OXA-48_, and bla ^3–5^. In the absence of carbapenemases, the loss of or defects in porins combined with an increase in *β*-lactamase expression can also mediate carbapenem resistance ^3–6^. However, the underlying mechanism of many CRKp strains remains undetermined, with some studies showing more than 50% of strains lacking a known functional carbapenemase^7,8^. Furthermore, our understanding of the genetic factors that facilitate the evolution of carbapenem resistance in clinical isolates is less well understood, including both intrinsic factors such as the genomic background of an isolate as well as extrinsic factors such as clinical antibiotic choice^6,9^.

Genome-wide association studies (GWAS) have successfully identified resistance-associated variants in other pathogens, including *Mycobacterium tuberculosis*, *Neisseria gonorrhoeae*, and *Staphylococcus aureus*^10^. These studies have revealed that known resistance genes often explain only a fraction of observed resistance^11^, while also uncovering candidate loci for point-of-care diagnostics^12,13^. In *K. pneumoniae*, three GWAS studies investigating carbapenem resistance have implicated biofilm formation^14^, capsular types and genes other than *β*-lactams associated with resistance^15,16^. While foundational, the existing studies in *K. pneumoniae* remain restricted in scope and methodology^17,18^. Each study examines clinical isolates only from a single country (India, China, or Italy) and does not cover a comprehensive set of genomic variants, *i.e.*, they test only genes or single nucleotide polymorphisms (SNPs) and do not accommodate other complex polymorphisms such as insertions, deletions and mobile genetic elements, which play a major role in the acquisition and spread of antimicrobial resistance^19^. Moreover, they do not account for the confounding effects of genome-wide linkage disequilibrium (LD)—the non-random association of alleles at two or more loci—which is a major contributor to strong population structure in bacteria because it leads to non-causal genomic elements hitchhiking with causal genomic variants, resulting in type Ι errors (false positives). Additionally, they employ a binary phenotype for modelling resistance, i.e., labelling strains as either resistant or susceptible instead of using a continuous value, which limits their ability to capture genetic variants with subtler contributions^15,20^.

To address these limitations, we conducted a GWAS on a set of *K. pneumoniae* clinical isolates from North America and Europe (spanning 33 countries), testing unitigs—variable-length nucleotide sequences derived from *de novo* genome assemblies—that represent a comprehensive set of genetic variants from the set of strains^21,22^. In addition, to increase statistical power to detect variants with subtle effects, we model the phenotype (resistance to meropenem), as a continuous trait using quantitative MIC values rather than a binary classification of resistant or susceptible. Furthermore, we integrated multiple analytical tools by employing an unweighted elastic net regression in conjunction with a linear mixed model (LMM), which has been previously proposed by Lees et al. and validated in *Mycobacterium tuberculosis*, *Neisseria gonorrhoeae*, *Streptococcus pneumoniae*, and *Streptococcus pyogenes* ^23,24^. Building upon the previous approach, we incorporated an additional variable in the LMM to control for the presence of carbapenemase genes known to confer resistance to meropenem. While the importance of this has been previously demonstrated by Ma et al for bacterial GWAS conducted using standard LMMs, combining it with an elastic net regression and LMM framework illustrates an extension of their work^25^. Although the underlying tools have been individually established, their combination has not been applied to *K. pneumoniae*. This integrated approach effectively controls for broad genetic confounders and filters out dominant signals of known resistance genes, yielding higher statistical power and sensitivity to isolate and prioritize genuinely novel genetic variants with independent effects on meropenem resistance.

Finally, to maximize the utility of these findings, we developed a visualization tool that integrates GWAS results, phenotypic data, and polymorphic site information. We use this tool to generate hypotheses for possible mechanisms by which a genetic variant could cause resistance, by examining the type of polymorphism and its relationship to meropenem resistance. The tool also aids in selecting strains for future experimental validation of the hypotheses.

Here, we show that by conditioning out carbapenemase genes, our method can more sensitively reveal genetic variants associated with both meropenem resistance and susceptibility. Strikingly, the majority of loci identified using this approach are mechanistically associated with either mutation or horizontal gene transfer (transposition, conjugation and phage infection), suggesting more broadly that *K. pneumoniae’s* genome plasticity itself underpins the acquisition of meropenem resistance.

## RESULTS

We collected 1,848 publicly available genome assemblies of *K pneumoniae* clinical isolates obtained from 248 hospitals, representing 33 countries in North America and Europe^21,22^. 1,705 of the 1,848 isolates had corresponding meropenem MICs and were thus utilized to conduct a bacterial GWAS (Fig. 1a). We converted the observed MIC values into discrete ranks to enhance the statistical power to detect genetic variants with smaller effects on the phenotype and also to ensure that the phenotypic data conformed to the underlying assumption of homoscedasticity required by the linear models employed as a part of the GWAS^20,25,26^. Then, to comprehensively represent the full extent of genomic variation present in our set of 1,705 strains, we extracted a total 5,025,653 unitigs from the genome assemblies of these isolates^27^ (see Methods). Unitigs are variable length nucleotide sequences that capture a wide range of genomic variants including SNPs, insertions, deletions, and mobile genetic elements across both the core and accessory genome without need for a reference genome^27^.

**Fig 1.**
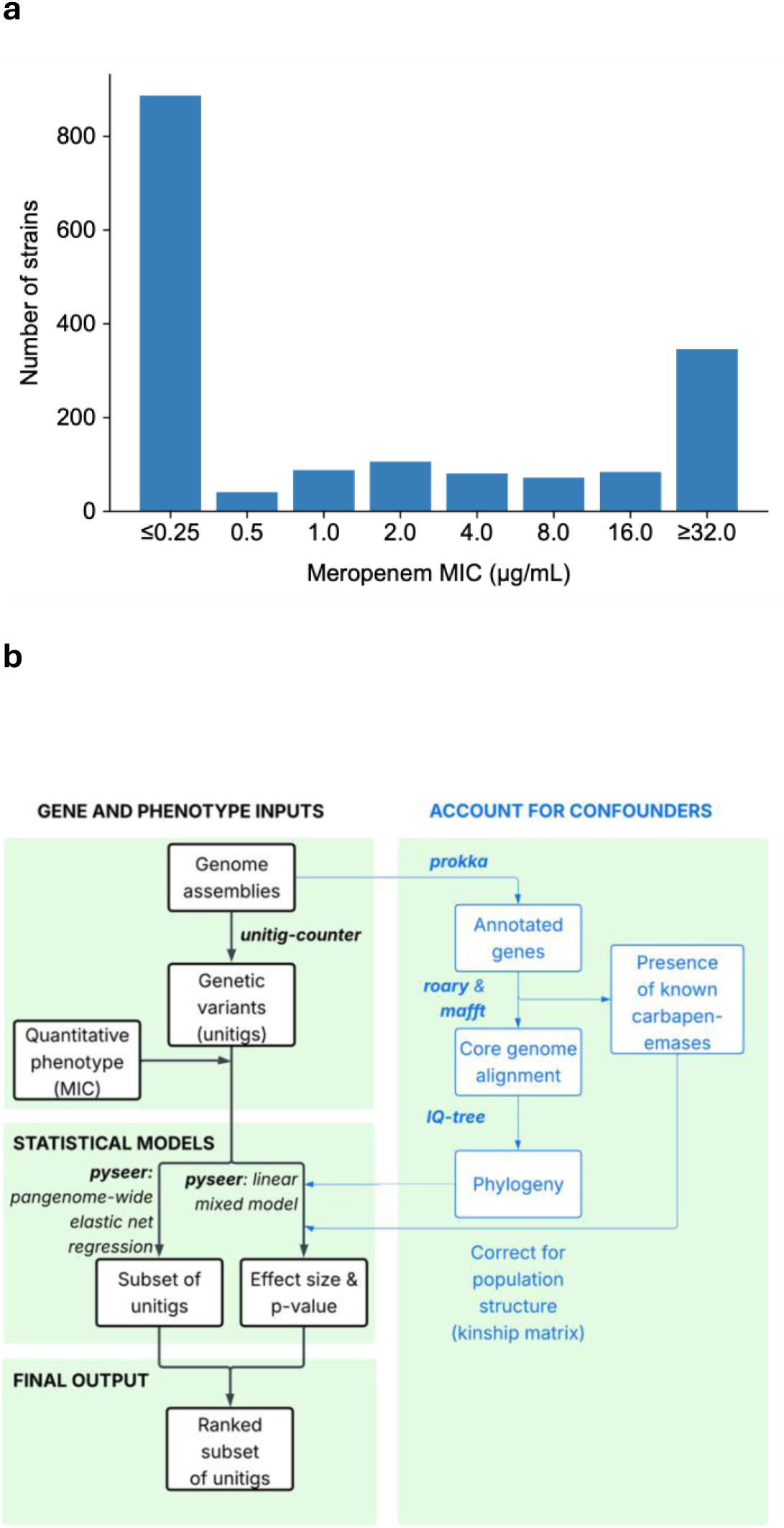
Phenotypic data and overview of GWAS workflow. **(a)** Distribution of the MIC values for the 1,705 *Klebsiella pneumoniae* clinical isolates used in the GWAS. MIC values are converted into discrete ranks of 1-8 before testing for association with the genotype. **(b)** Workflow used for conducting the GWAS. We use two different statistical methods: 1) an elastic net regression, which controls for pangenome-wide linkage by simultaneously testing all variants (unitigs), 2) a conditional linear mixed model, which not only corrects for population structure using a kinship matrix, but controls for the presence of carbapenemase genes known to confer resistance. The output of these two methods are then combined to obtain a subset of genomic variants associated with the phenotype.

To account for confounding factors such as slow decay of LD and clonal reproduction which lead to strong population structure^23,29–31,32^, we utilized both a multi-locus (*i.e.*, testing multiple genetic variants simultaneously) elastic net regression and a single-locus (*i.e.*, testing each genetic variant independently) LMM conditioned on known carbapenemase genes (Fig. 1b). The multi-locus elastic net implicitly includes interactions and correlations between unitigs and excludes those that co-segregate due to population structure and are not predicted to affect resistance^23,33^, thereby accounting for pangenome-wide LD and improving power to detect variants strongly correlated with lineage compared to using a LMM^23,30^ (Fig. 2). Moreover, by including known carbapenemase genes as covariates in the conditional LMM, we downweight confounding variants that co-segregate with these genes, resulting in increased power to detect variants with independent and subtler effects on meropenem resistance compared to using an unconditional LMM^23,25,34,35^ (Fig. 2).

**Fig 2.**
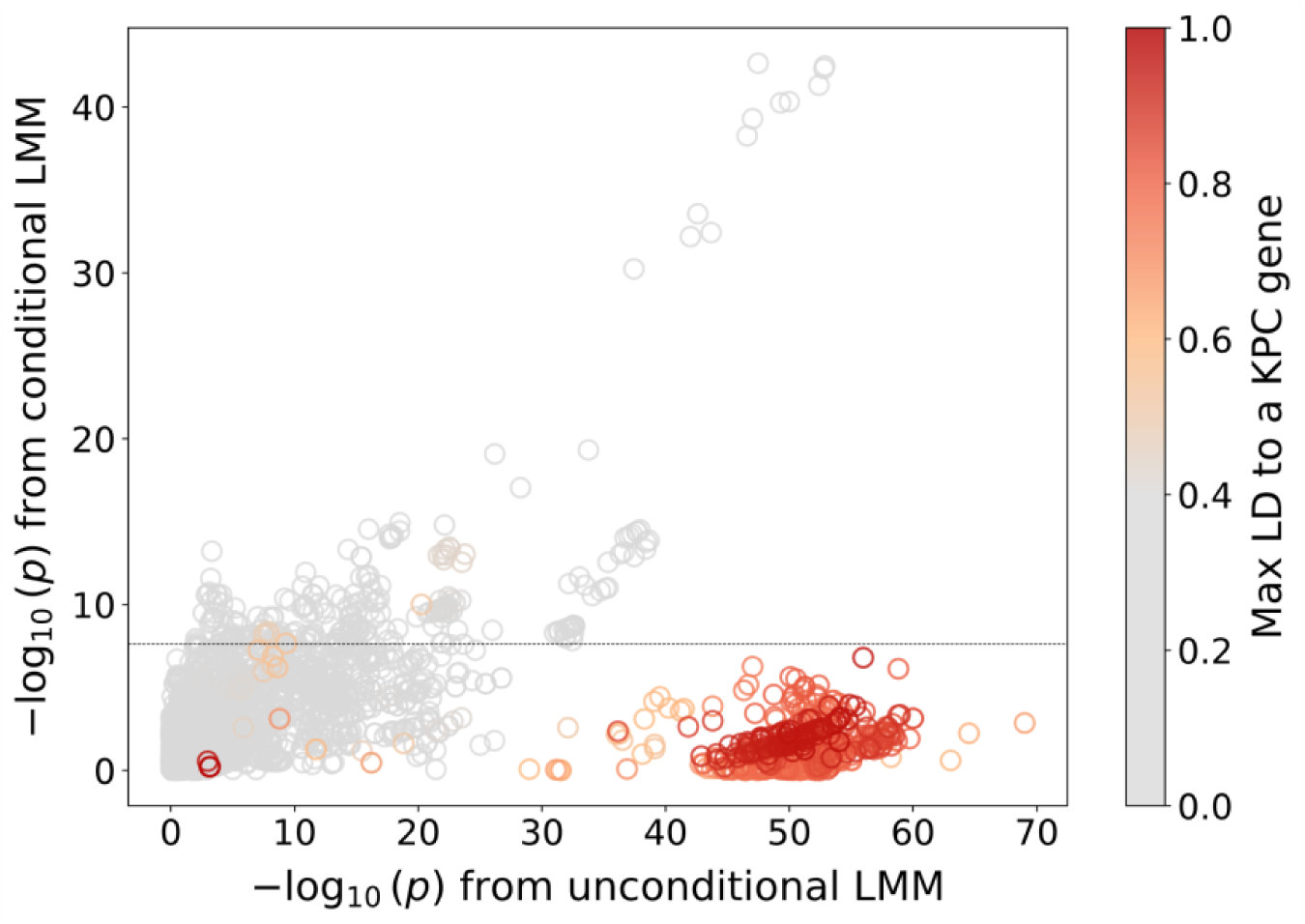
Conditional GWAS accounting for known carbapenemase genes reduces confounding by reducing the statistical significance of variants that co-segregate with them. The negative log_10_-transformed *p*-values obtained from the unconditional and conditional linear mixed model (LMM) are plotted for each of the 3,836 unitigs selected by the elastic net regression. Linkage disequilibrium (LD) measured by *r*^2^ to a set of 12 carbapenemase orthologs is colored for all variants, ranging from 0 (light grey) to 1.0 (dark red). A Bonferroni threshold calculated using the number of unique patterns for the conditional GWAS is depicted using a dashed grey line at 7.67 x 10^7^.

Thus, we performed a GWAS on the final set of 1,705 strains by applying both an unweighted multi-locus elastic net^23,36^ and a single-locus conditional LMM. In both models, we tested 2,358,722 unitigs that had allele frequencies in between 1-99% (*i.e.,* present in 1-99% of strains) against the associated ranked meropenem MICs. The elastic net with a penalty of α = 0.01 yielded a set of 3,836 unitigs (Supplementary Table 1) predicted to be associated with the phenotype. We chose α = 0.01 for the elastic net penalty to reduce sparsity and lower the likelihood of excluding causal variants from our final set^23^. We then ranked these unitigs from the elastic net by their statistical significance determined independently for each unitig from the conditional LMM.

We observed that the 3,836 unitigs represented distinct types of genetic variants including SNPs, insertions, allelic variants, and mobile genetic elements, mapping to 306 genes and 149 intergenic regions. We focused on eight loci (Fig. 3) implicated by 125 of the 300 most statistically significant unitigs of which 80 had a positive effect size, i.e., were associated with higher MIC values or increased meropenem resistance, and 45 had a negative size, i.e., were associated with lower MIC values or increased meropenem susceptibility (Table 1). The remaining 175 unitigs of the top 300 were annotated as lying in putative hypothetical proteins, genes encoding domains of unknown function, or intergenic regions. Hence, functional mechanisms by which these latter categories of unitigs might be associated with resistance or susceptibility to meropenem were more difficult to predict and they were therefore filtered out from further analysis (Supplementary Table 2).

**Fig 3.**
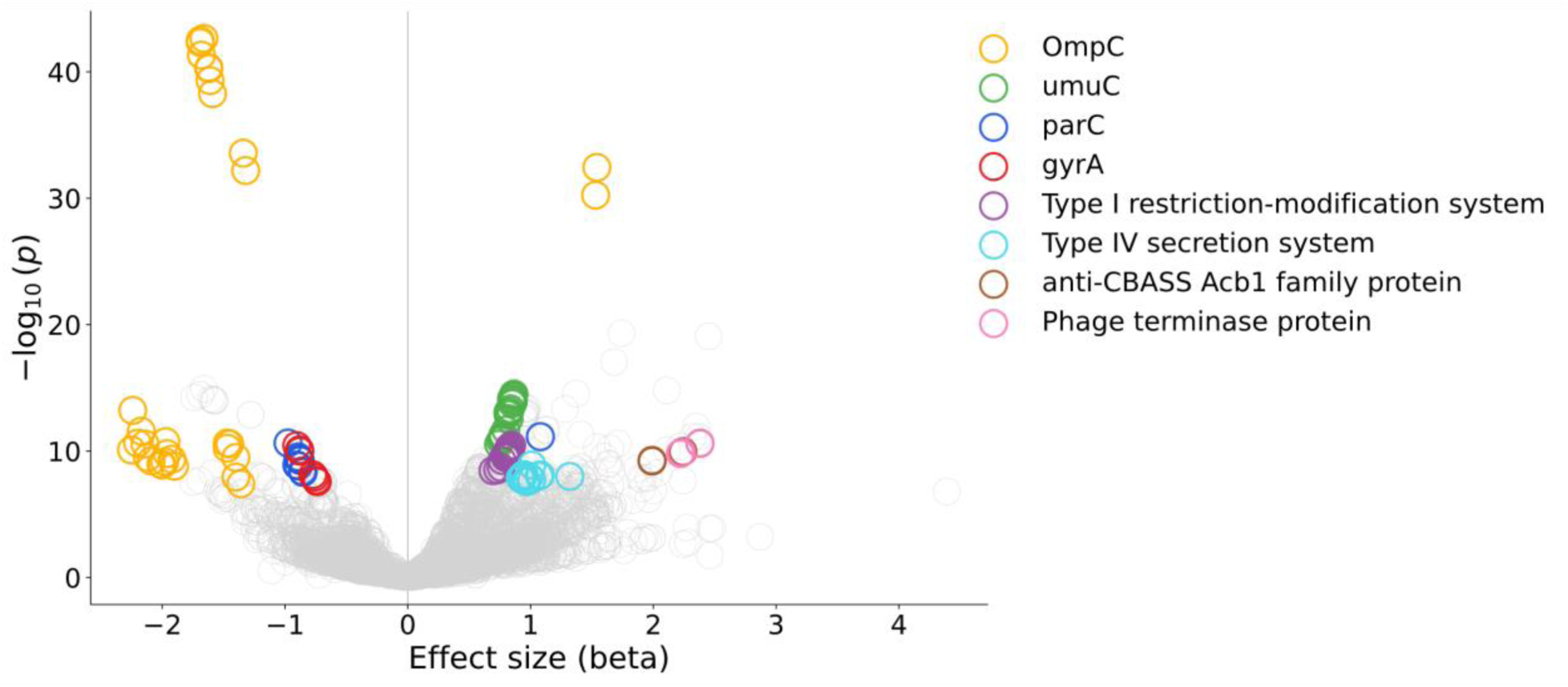
Volcano plot highlights unitigs corresponding to the eight candidate loci from the GWAS. A total of 3,836 unitigs are selected by the elastic net regression and plotted using their negative log_10_-transformed *p*-value from the conditional linear mixed model. Unitigs with a positive beta value are predicted to be associated with meropenem resistance, and unitigs with a negative beta value are predicted to be associated with meropenem susceptibility.

**Table 1.** A list of the eight candidate loci representing 125 of the 300 most statistically significant unitigs (of a total of 3,836) selected by the elastic net.

| **Candidate locus** | **Variant type** | **Number of top 300 unitigs from the elastic net mapping to it** |  | **P-value range (Min, Max)** |  |
|---|---|---|---|---|---|
|  |  | **Resistance-associated (mean beta)** | **Susceptibility-associated (mean beta)** | **Resistance-associated** | **Susceptibility-associated** |
| *ompK36* | Insertion | 2 (1.54) | 30 (-1.78) | 3.68E-33, 5,73E-31 | 2.28E-43, 6.49E-33 |
| *umuC* | Allelic variant | 20 (0.82) | 0 | 3.09E-15, 2.86E-11 | - |
| *hsdRMS-IsKpn25* | Mobile genetic element | 22 (0.80) | 0 | 3.39E-11, 3.51E-09 | - |
| Type IV secretion system (*virB9, virB10, virB11* ) | Allelic variant | 20 (0.98) | 0 | 1.25E-09, 2.62E-08 | - |
| *parC* | SNP | 1 (1.08) | 9 (-0.69) | 7.19E-12 | 2.28E-11, 4.85E-09 |
| *gyrA* | SNP | 0 | 6 (-0.81) | - | 3.66E-11, 2.63E-08 |
| anti-CBASS *Acb1* | Allelic variant | 10 (2.26) | 0 | 2.40E-11, 6.13E-10 | - |
| Phage terminase protein | Allelic variant | 4 (2.3) | 0 | 2.53E-11, 1.58E-10 | - |

The first of the eight loci corresponded to the *ompK36* gene which encodes the major outer membrane porin OmpK36^37^. The 12 most significant unitigs all mapped to this gene, with two of them having positive effect sizes (*β* = 1.54 and 1.53, *p-*values = 3.68 × 10^-33^ and 5.73 × 10^-31^, respectively) and the remaining ten with negative effect sizes (mean *β* = -1.58, least significant *p-*value = 6.49 × 10^-33^). The unitigs associated with negative effect sizes, and therefore meropenem susceptibility, corresponded to the wild-type *ompK36* allele, which is permissive for meropenem entry. In contrast, one of the of significant unitigs with positive effect size (*β* = 1.53; meropenem resistance) corresponded to a 6-bp Gly-Asp (GD) insertion *134_135insGD* known to increase resistance to carbapenems by constricting the pore and restricting diffusion of carbapenems across membranes^38–40^. The other significant unitig with a positive effect size (*β* = 1.54) corresponded to T136A substitution in the L3 loop of the *ompK36* β-barrel. While it is possible that this conservative substitution might affect pore size, it is in strong linkage disequilibrium (*r*^2^ = 0.95) with the *134_135insGD* insertion and hence its significance might be almost entirely attributed to the known *134_135insGD* insertion.

Surprisingly, the second and third loci corresponded to the *parC* and *gyrA* genes, which encode the catalytic subunits of Topoisomerase IV and DNA gyrase, respectively, and are the primary targets of fluoroquinolone antibiotics, not carbapenems. Eight unitigs mapped to *gyrA,* seven of which had a negative effect size (mean *β* = −0.81, least significant *p-*value = 2.63 × 10^-8^) and one of which had a non-significant positive effect size because of its low allele frequency of 0.01 (i.e., present in only 1% of strains). Ten unitigs mapped to *parC*, with all but one unitig having a negative effect size (*mean β* = −0.69, least significant *p-*value = 4.85 × 10^-9^), similar to *gyrA*. The single *parC* unitig with positive effect size (*β* = 1.08, *p-*value = 7.19 × 10^-12^) mapped to an S80I substitution known to confer resistance to fluoroquinolones.

All unitigs with negative effect sizes in *gyrA* overlapped one or both of the most frequently mutated residues (Ser83 and Asp 87) in the quinolone resistance determining region (QRDR) that confer resistance to fluoroquinolone antibiotics. Likewise, all 9 *parC* unitigs with negative effect sizes mapped to one of the most frequently mutated residues in *parC* that confers fluoroquinolone resistance (Ser80). In both genes, the vast majority of unitigs are precise matches for fluoroquinolone susceptible alleles of *gyrA* and *parC* and given their negative effect sizes, are also associated with susceptibility to meropenem. Rather than conferring meropenem susceptibility by some novel mechanism, we postulate that the fluoroquinolone susceptible alleles marked by these unitigs are associated with meropenem susceptibility because these traits are tightly linked in evolutionary time and reflect the relative short time frame in which strains are exposed to these antibiotics clinically. Hence, we predict that meropenem susceptible stains are more likely to be fluoroquinolone susceptible and conversely, meropenem resistance strains are more likely to be fluoroquinolone resistant.

The fourth locus corresponded to the *umuC* gene, which encodes a catalytic subunit of a Y-Family DNA polymerase V and with *umuD_2_*is involved in SOS-induced error prone repair^41,42^. It has also been implicated in carbapenem resistance in a previous genome-wide association study^15^, but has not been functionally or experimentally validated^14–16^. We observed 20 unitigs that mapped to this gene, which all had positive effect sizes and were statistically significant (mean *β* = 0.818, least significant *p-*value = 2.86 × 10^-11^). We observed that the unitigs overlap to cover a large part of the *umuC* gene from codons 169-343, likely indicating that a specific *umuC* allele is associated with meropenem resistance. We hypothesized that Y-family translesion synthesis (TLS) polymerase induced as part of the SOS response by *umuC* enables replication past unrepaired lesions due to chemical or DNA-induced damage, leading to stepping-stone mutations that result in increased resistance to meropenem^43,44^. It also remains a possibility that the specific allele of *umuC* mapped by the 20 unitigs confers strain-specific high-level resistance to meropenem by a novel mechanism.

For the fifth locus, we observed 22 significant unitigs associated with increased meropenem resistance (mean *β* = 0.799, least significant *p-*value = 3.51 × 10^-9^) that map to the *hsdM-hsdS-hsdR-IsKpn25* genes, tiling across the entire element. The *hsdM-hsdS-hsdR* (*hsdMSR*) genes make up a type I restriction modification system that is frequently found in mobile elements of the genome such as plasmids, phages, and transposons^45^. The *IsKpn25* gene is an insertion sequence (IS) element belonging to the ISL3 transposase family which has previously been shown to mediate colistin resistance in CRKp by inserting itself and the *hsdMSR* genes into the *mgrB* gene, inactivating it in the process^45,46^.

We observed the unitigs that map to the *hsdM-hsdS-hsdR-IsKpn25* element mapped to two distinct loci in the MGH_78578 reference strain^47^; one in the pKPN4 plasmid and the other in the chromosome, adjacent to the *ompK36* gene. Given the presence of target site duplications immediately flanking the inverted repeats at both loci, the *hsdM-hsdS-hsdR-IsKpn25* element integrated at these sites via transposition, indicating that it is an active transposase. This element could confer resistance to meropenem by increasing mutation frequency via transposition, in the process leading to frequent and reversible disruption of porin genes^6^. However, we could not find evidence of transposition into different regions of the genome in the set of 288 strains the element was present in, possibly because of the high level of fragmentation in the genome assemblies of the strains, so the degree to which IsKpn25 transposition contributes to meropenem resistance is undetermined. The *hsdM-hsdS-hsdR-IsKpn25* element is also known to be carried by pKpQIL,^48,49^ which encodes the carbapenemase gene *bla*_KPC-3_ and is frequently identified in CRKp lineages such as ST512 and ST258^45,50^. Alternatively, or in addition to transposition, another possibility is that the *IsKpn25* gene has co-opted the *hsdMSR* restriction modification system to protect itself from degradation where it jumps and may function to protect plasmids that encode it and cells where it is harbored from post-segregational killing^51^. It may therefore be associated with meropenem resistance by ensuring plasmid propagation.

We noticed that the *hsdMSR* genes co-segregate frequently with the *umuC* allele associated with resistance (Fig. 4), raising the concern that one of them was a spurious association due high LD with the other. To test this, we conducted two statistical tests, each employing a conditional LMM for determining whether *umuC* or the *hsdM-hsdS-hsdR-IsKpn25* element was a confounder. The tests indicated that both umuC and the *hsdM-hsdS-hsdR-IsKpn25* element have independent effects on the meropenem MIC (Supplementary Information). We also observed that the individual effects of umuC and the *hsdM-hsdS-hsdR-IsKpn25* element were not linearly additive, postulating this could be explained by redundance in the biological mechanisms by which each confers resistance or the saturation of the assay used to measure resistance itself.

**Fig 4.**
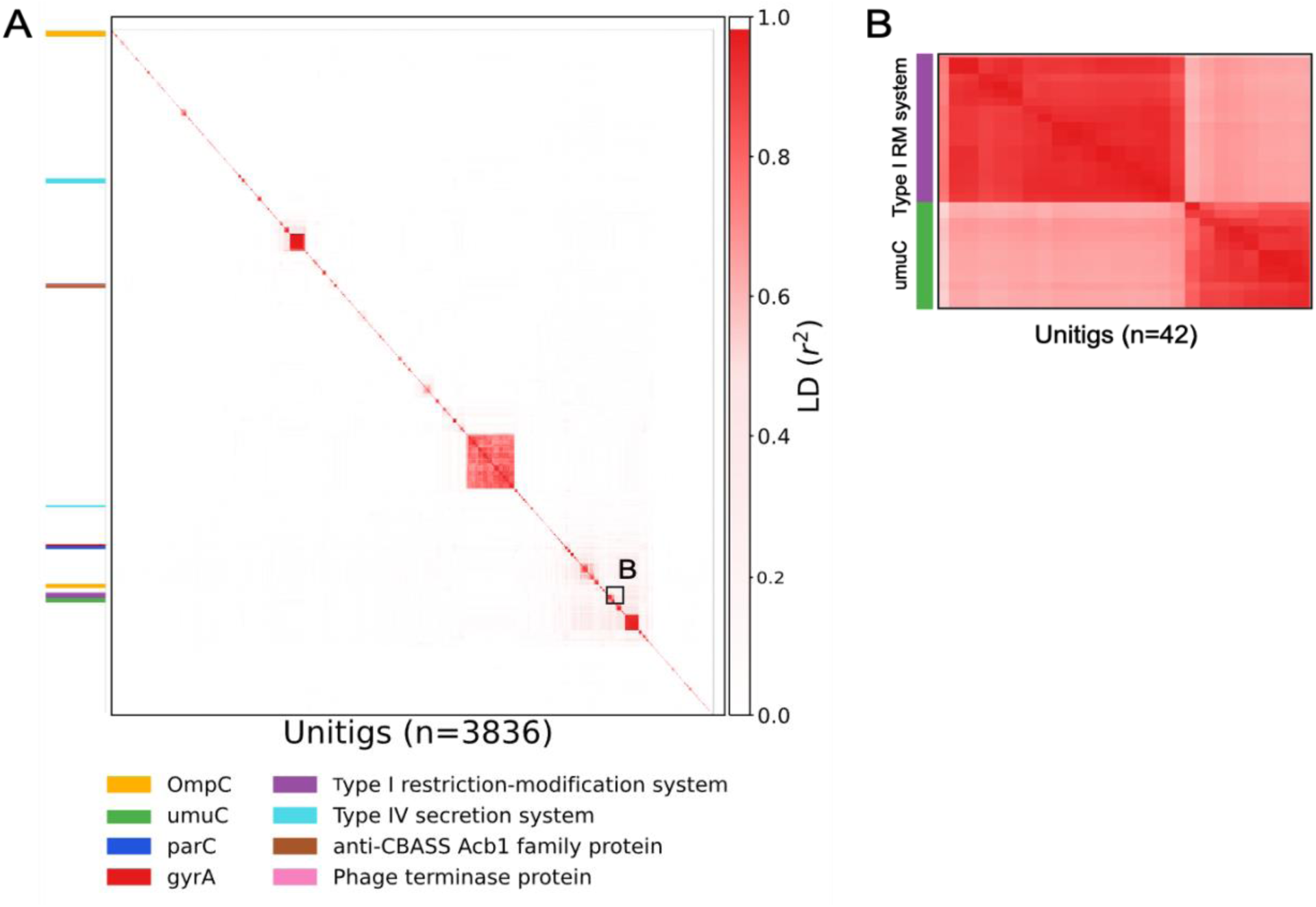
Heatmap illustrating linkage disequilibrium between unitigs selected from the elastic net regression. Linkage disequilibrium (LD) is measured by calculating *r*^2^ between each pair of the 3,836 unitigs. The heatmap portrays distance-based clustering of these unitigs and shows that *umuC* and *hsdRMS-IsKpn25* co-segregate with each other frequently (panel **A**), while the other six candidates map to separate blocks.

At the sixth locus, we observed 13 unitigs mapping to the *virB11*, *virB10*, and *virB9* genes, which are components of a Type IV secretion system (T4SS) that enable conjugative transfer of plasmids between bacterial cells^52,53^. Six unitigs mapped to *virB11* (mean *β* = 0.968, least significant *p-*value = 1.68 × 10^-8^), five unitigs to *virB10* (mean *β* = 0.979, least significant *p-* value = 2.62 × 10^-8^), and two unitigs to *virB9* (mean *β* = 1.075, least significant *p-*value = 9.44 × 10^-9^) We conjectured that the resistance-associated unitigs correspond to a T4SS system that promotes greater conjugation frequency, leading to increased resistance to meropenem.

The seventh and eight loci we observed corresponded to an anti-CBASS *Acb1* gene and Phage terminase protein located immediately adjacent to one another, respectively. Ten unitigs mapped to the anti-CBASS *Acb1* gene (mean *β* = 2.259, least significant *p-*value = 6.13 × 10^-10^) and four unitigs that map to a Phage terminase (mean *β* = 2.3, least significant *p-*value = 1.58 × 10^-10^). We hypothesized that *Acb1* renders cells more susceptible to phage infection, thereby creating a more permissive environment for horizontal gene transfer and possibly leading to the acquisition of genes that lead to increased resistance to meropenem^6,54^ and that the unitigs corresponding to the Phage terminase protein were identified based on their linkage to *acb1*.

Last, to aid in developing hypotheses concerning the role of the candidate loci in meropenem resistance and to guide future experimental design to confirm them, we developed a new visualization tool. This tool integrates the GWAS results, phenotypic data, and polymorphic site information, and allows selection of a specific subset of clinical isolates for further experimental validation (Fig. 5, see Methods). We observed that for u*muC, hsdM-hsdS-hsdR-IsKpn25,* the *virB9, virB10, and virB11* genes comprising of the T4SS system, anti-CBASS *Acb1* gene and Phage terminase protein, the mapped unitigs overlap or “tile” a significant portion of the genes, indicating a particular allelic variant of the gene is associated with the phenotype. Meanwhile, the unitigs for *ompK36, parC* and *gyrA* are stacked under a single codon and encode a single SNP (Fig. 5). This information guided our hypotheses that specific allelic variants involved in mutagenesis, transpositional insertion, conjugal transfer, and phage infection in *Klebsiella pneumoniae* lead to the acquisition of meropenem resistance.

**Fig 5.**
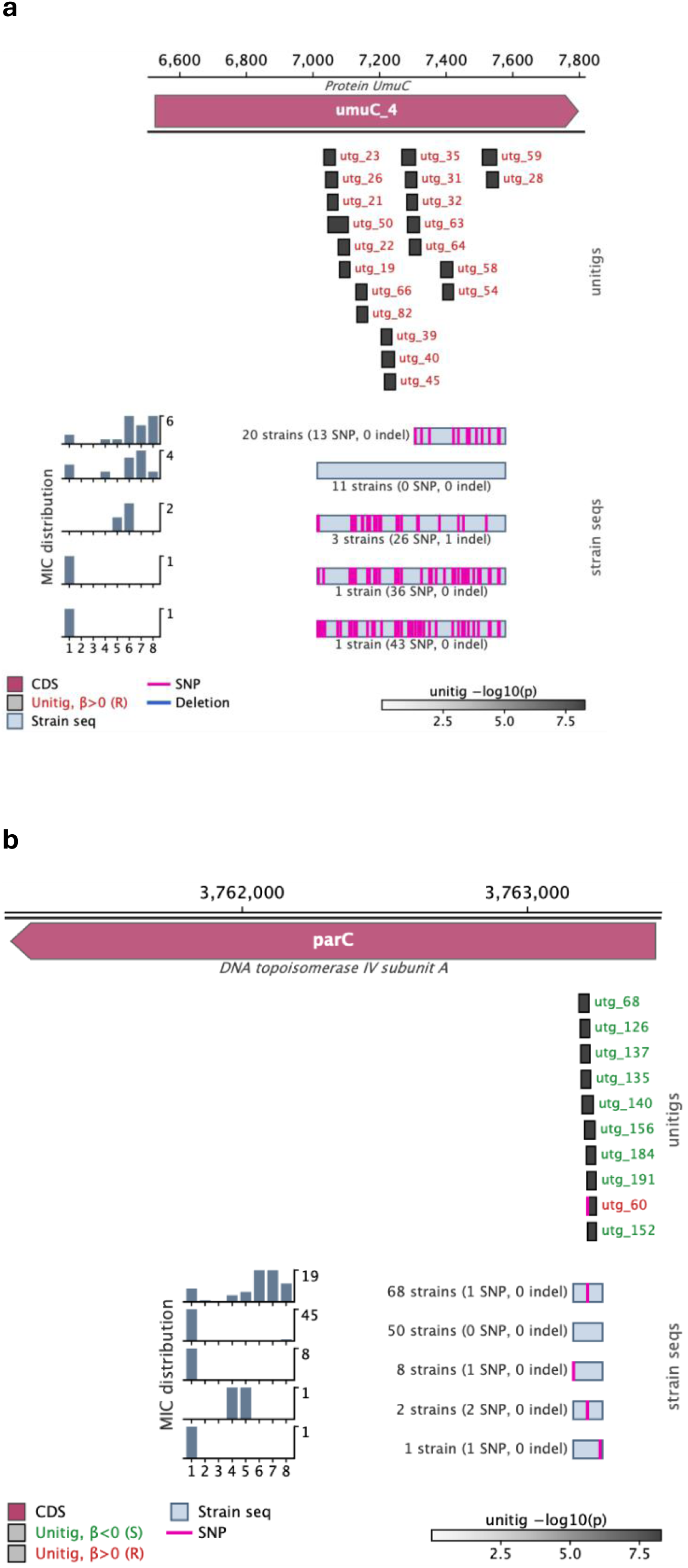
Examples of visualization tool, integrating GWAS results with polymorphic site information from unitigs. **(a)** We observe 20 statistically significant unitigs that ‘tile’ or overlap a significant section of *umuC* in the BIDMC_53 strain, from which we infer an allelic variant of *umuC* is associated with meropenem resistance. By aligning these unitigs to the subset of 131 strains from Cerqueira et al. used in our GWAS, we observe that 11 strains from this subset have the same sequence of *umuC* as in BIDMC_53. **(b)** We observe 10 statistically significant unitigs that stack under a single codon of *parC* in the MGH_78578 reference strain, with one encoding a S80I substitution known to confer resistance to fluoroquinolones. From this we infer a single SNP in the *parC* is associated with meropenem resistance. By aligning these unitigs to the subset of 131 strains from Cerqueira et al. used in our GWAS, we observe that 68 strains from this subset encode the S80I substitution. For each panel, we select a total of 5 sequences extracted from the 131 strains to display, sorting by sequences most commonly observed. If two sequences have the same strain count, we display the one with a lower median MIC across the strains it is present in.

## DISCUSSION

Conventional approaches in bacterial GWAS studies for uncovering the unknown genetic basis of antimicrobial resistance have encountered challenges in detecting novel genomic variants associated with lower effect sizes that play a critical role in the acquisition and spread of resistance^25^. By incorporating known resistance genes (conditional LMM) and accounting for LD spanning the entire genome (elastic net regression) in our GWAS, we were able to obtain the statistical power to detect novel genomic variants with smaller effect sizes that represent a broad set of polymorphisms including insertions, deletions, and mobile genetic elements. By using a conditional instead of an unconditional LMM, we were able to demonstrate a decrease in the number of spurious associations attributable to genetic linkage with KPC genes.

By developing a new visualization tool that integrated information from our GWAS results, phenotypic data, and polymorphic site information across our set of strains we were able to construct hypotheses for a majority of the 306 unique genes that were mapped by the unitigs selected by the elastic net regression. We identified two distinct patterns across the visualizations we generated, i.e., cases where the unitigs overlapped significant sections or “tiled” a gene or multiple genes at a locus and other cases where they stacked under a single codon and encoded a mutation such as SNP or insertion.

By analyzing these visualizations, we observed that many of the most statistically significant unitigs tiled novel genes such as *umuC,* the *hsdMSR* genes comprising of the type I restriction modification system, as well as the *virB9*, *virB10*, and *virB11* genes that form part of a T4SS system. From these observations that large sections of these genes were overlapped by several unitigs from our GWAS, we hypothesize that specific allelic variants (rather than individual SNPs) could influence the propensity to evolve resistance via facilitating stepping-stone mutations that confer resistance, transpositional mutagenesis, and increasing conjugation frequency.

Comparing our results to existing GWAS studies performed in *K. Pneumoniae* that investigated carbapenem resistance, we observed that Pei et al. was the only one to focus on uncovering mechanisms associated with meropenem resistance^14–16^. Similar to the results of the GWAS conducted in this study, Pei et al. also established an allele of *umuC* and the Ser83 mutation in *gyrA* to be associated with meropenem resistance in their GWAS. However, we cannot determine if the *umuC* allele they identified matches the variant we selected because the sequence of their resistance-associated ortholog was not reported. All other 11 genes and 13 SNPs computationally validated by Pei et al. are not selected by our GWAS as statistically significant. Meanwhile, none of the loci mapped by the 300 most statistically significant unitigs from our GWAS appear in Pei et al.’s results. This could be due to differences in geographical origin of the clinical isolates or discrepancies in variant annotation. In addition, our GWAS prioritized variants with low LD to carbapenemase genes because of conditioning which could further explain the observed lack of overlap.

To determine whether any of the eight candidate loci we prioritized could explain resistance in strains without known determinants of carbapenem resistance, we analyzed the 30 strains of unexplained resistance (*i.e*., strains that have an MIC of ≥ 1 μg/mL and do not have any known genetic factors that confer carbapenem resistance) in the set of 1,705 strains used in our GWAS^5,21,22^. We examined whether any of five of the eight candidate loci — *umuC, hsdM-hsdS-hsdR-IsKpn25,* the *virB9, virB10, and virB11* genes comprising of the T4SS system, anti-CBASS *Acb1* gene and Phage terminase protein — were present in the strains of unexplained resistance. Of the 30 clinical isolates of unexplained resistance, we observed that ten have the allelic variant of *umuC* present, nine have the *hsdM-hsdS-hsdR-IsKpn25* present, five have the T4SS system with the allelic variants of *virB9, virB10, and virB11* present, and none have either the anti-CBASS *Acb1* gene or Phage terminase protein. Eight strains of unexplained resistance have more than one of these five candidate loci present.

Experimental validation is required to confirm the mechanisms by which the candidate loci selected by our GWAS are conjectured to induce or confer increased resistance to meropenem. Selecting strains of unexplained resistance with the relevant genetic background with the help of the visualizations we produced could be a first step in designing an experiment to test these candidates.

Our study is not without its limitations. Although we reduced type Ι error (false positives) through performing a conditional LMM, we acknowledge that using an elastic net penalty of α = 0.01 as opposed to conducting a lasso regression (α = 1) risks higher type Ι error. However, using a lasso regression would lead to reduced power to detect variants with smaller effect sizes^23,30^. It is also difficult to generate hypotheses for unitigs that mapped to intergenic regions. The effect of these mutations could be explored by integrating GWAS results with eQTL studies to determine which of these intergenic regions are associated with statistically significant changes in gene expression and in which genes.

In summary, by reducing the effect of genomic confounders such as population structure due to genome-wide LD and clonal reproduction through combining the results of an elastic net regression with those of a LMM conditioned on known genes, we were able to identify novel genetic variants associated with meropenem resistance. Furthermore, via the visualization tool we developed, we produced hypotheses for a set of novel candidates and prioritized candidates and strains to guide future experiments. Experimental validation of these candidates along with integrating GWAS results with eQTL studies are important avenues for future research. The results from the GWAS illustrate that mechanisms underpinning genome plasticity in *Klebsiella pneumoniae* likely play an important role in the acquisition of meropenem resistance.

## METHODS

### Data Collection and QC

Genome assemblies and the corresponding MIC measurements for each *Klebsiella pneumoniae* clinical isolate used in this study were downloaded from NCBI BioProject accession number PRJNA202876^21^ (n = 131) and European Nucleotide Archive (ENA) study accession no. PRJEB10018^22^ (n=1717). 1,574 of the isolates from the ENA study along with the 131 *Klebsiella pneumoniae* isolates from the PRJNA202876 BioProject had corresponding meropenem MIC values and together were selected for the GWAS. The meropenem MICs of these 1,705 genome assemblies were converted into discrete ranks ranging from 1 to 8.

Kleborate v3.2.4^55,56^ was used for quality control of these genome assemblies with meropenem MIC values. Assemblies with N50 < 10 kb, total size > 7.5 Mb or < 4.5 Mb, or with ≥ 3% of their total length consisting of ambiguous bases were discarded.

Assembly accessions and quality control statistics for the strains used in the GWAS are available in Supplementary Table 3.

### Gene Annotation and Phylogeny Construction

All 1,848 genome assemblies were annotated with Prokka v1.14.6^57^. The annotated assemblies were used as input for Roary v3.12.0^58^ (flags -e -n -v -p 8 -i 90) to build a pangenome and determine the core and accessory genes across the isolates. The pangenome comprised of 56,014 genes in total, including 2,989 core genes (*i.e.,* genes present in ≥ 99% of strains). An alignment of the 2,989 core genes was generated using Mafft v7.508^59^. Because of the high genomic diversity in our set of strains, removing recombinant regions from the core gene alignment before generating a phylogeny was not feasible^22,60^. Therefore, we extracted the variable positions from the core gene alignment and generated a maximum likelihood tree to represent the phylogeny of the isolates using IQ-TREE v2.0.3^61^ with a general time reversible model (flags -fast -m GTR).

We used the phylogeny to construct a kinship matrix *K* that describes the similarity between isolates using the phylogeny_distance.py script from pyseer v1.3.10 with the --lmm flag^24^.

### Identification of Known Resistance Alleles

Carbapenemase genes were detected using a custom database with known resistance genes (Supplementary File 1). The pan_genome_reference.fa file containing one representative nucleotide sequence from each gene cluster in the pangenome was used as a query file to run BLASTn v2.16.0^62^ on the custom database. We identified a gene as a carbapenemase if it had ≥ 90% identity and ≥ 80% coverage of the resistance allele’s length. We identified 12 carbapenemase orthologs belonging to one of the five major carbapenemase families of bla_KPC_, bla_NDM_, bla_OXA-48_, bla_VIM_, or bla_IMP_.

### GWAS

We extracted unitigs from the 1,705 *Klebsiella pneumoniae* genome assemblies using unitig-counter^27^. We tested the association of each unitig present in 1-99% of strains with meropenem MIC independently, controlling for population structure and linkage to carbapenemase genes by applying a conditional linear mixed model using pyseer v1.3.10^24^ (flag --lmm)

#### Model

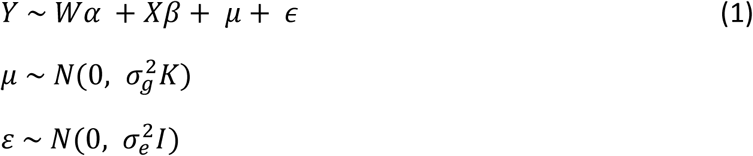

Here, *Y* is a vector of meropenem MICs that have been converted into discrete ranks of 1-8 for all the 1,705 strains used in the GWAS. *W* is the covariate matrix which includes for each strain the dataset of origin as well as the presence or absence of the 12 carbapenemase genes that are known resistance alleles. *α* represents the fixed effect for each covariate in *W*. *X* is a vector representing the presence of the unitig being tested, with *β* being its fixed effect (*i.e.,* effect on the meropenem MIC). *μ* is the random effect to control for population structure parametrized with the similarity matrix *K* calculated from the phylogeny and additive genetic variance 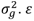 is the random effect that models non-genetic or environmental effects and is parametrized with variance 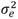 and identity matrix *I*.

We also selected for genetic variants that best predict the phenotype ( *i.e.,* meropenem MIC) by testing all unitigs simultaneously, controlling for pangenome-wide linkage disequilibrium by applying an elastic net regression using pyseer v1.3.10 (flags --enet --alpha 0.01 --n-folds 10 -- cor-filter 0)^23,24^.

#### Model

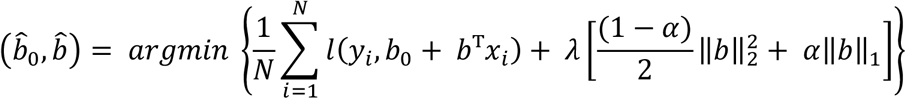

Here, *b*_0_ is the intercept and *b* is a vector representing the slopes or fixed effects for each unitig tested in the elastic regression. *N* is the number of samples (*i.e.*, 1,705) and *y_i_* represents the meropenem MIC of the *i^t^*^ℎ^ strain. *l* refers to the link function, which is linear for a continuous phenotype like MIC. *x_i_* is a vector representing the presence or absence for each unitig tested for the *i^t^*^ℎ^ strain. *λ* is the parameter used to control penalty strength that is chosen by cross-validation. *α* is the amount of mixing between the ridge (L1) and lasso (L2) penalty. *b̂*_0_ is the predicted intercept value and *b̂* is vector of the predicted fixed effect sizes for each unitig. The elastic net penalty shrinks most of the predicted effect sizes to zero, finally selecting the unitigs from *b̂* that are predicted to have non-zero effect sizes. We used *α* = 0.01 to reduce sparsity and control for false negatives, i.e., prevent excluding unitigs that affect meropenem MIC from those selected by the elastic net.

The fit of the elastic net is measured using *R*^2^ calculated using the formula:

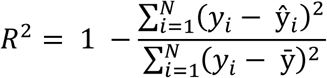

where ŷ*_i_* is the predicted meropenem MIC of the *i^t^*^ℎ^ strain using the elastic net.

By combining the results of the elastic regression with the conditional linear mixed model, we obtained a set of 3,836 unitigs (with *R*^2^ = 0.89) predicted to affect meropenem MIC from the elastic net that can be prioritized by their statistical significance obtained from the conditional linear mixed model.

### Linkage Disequilibrium Calculation

We calculate linkage disequilibrium (LD) for using the squared correlation coefficient

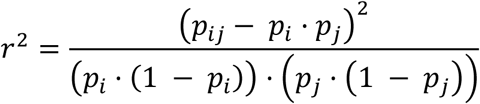

Where 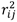 is the LD or squared correlation between unitigs *i* and *j*. *p_i_* is the proportion of strains with unitig *i*, *p_j_* is the proportion of strains with unitig *j*, and *p_ij_* is the proportion of strains with both unitigs *i* and *j*.

For each of the 3,836 unitigs selected by our GWAS, we calculated LD between each pair using *r*^2^. We also calculated the LD between each of the 3,836 unitigs and the 12 carbapenemase genes using *r*^2^, taking the maximum value to assess genetic linkage between a unitig and carbapenemase gene.

### Testing Independent Effects of *umuC* and *hsdRMS-IsKpn25*

For testing whether *umuC* affects the phenotype independent of the presence of *hsdRMS-IsKpn25*, we perform a conditional linear mixed model using pyseer (flag --lmm)^24^ similar to as described in (1). We modified the covariate matrix *W* so that it includes the presence and absence of the 22 resistance-associated unitigs mapping to *hsdRMS-IsKpn25*, in addition to the dataset of origin and the presence or absence of the 12 carbapenemase genes that are known resistance alleles. We then independently test each of the 20 resistance-associated unitigs mapping to *umuC* again using the conditional linear mixed model.

Similarly, for testing whether *hsdRMS-IsKpn25* affects the phenotype independent of the presence of *umuC*, we also perform a conditional linear mixed model using pyseer (flag -- lmm)^24^. We modified the covariate matrix *W* so that it includes the presence and absence of the 20 resistance-associated unitigs mapping to *umuC*, in addition to the dataset of origin and the presence or absence of the 12 carbapenemase genes that are known resistance alleles. We then independently test each of the 22 resistance-associated unitigs mapping to *hsdRMS-IsKpn25* again using the conditional linear mixed model. We also separately tested a single vector representing the presence and absence of the entire element *hsdRMS-IsKpn25* by applying a conditional linear mixed model using pyseer^24^ (Supplementary Information).

### Unitig Annotation

All 3,836 unitigs were mapped to the *Klebsiella pneumoniae* reference genome MGH_78578 using bwa-mem^63^ (with parameters -k 8 -B 2 -O 3) and to all 1,705 strains used in the GWAS using bwa fastmap^63^ (with parameters -l 9 -w 100). We developed scripts align_and_annotate.py and plot_unitigs_compact.py to annotate and visualize mapped unitigs. SnpEff v5.0^64^ was used to predict the functional effect of unitigs.

## Code Availability

Claude Code models Sonnet 5.5 and Opus 5.5 were used to assist with polishing code developed for the visualization tool created to generate hypotheses for genomic variants. We reviewed, modified and tested all code and validated its outputs by comparing its output with source data and validating the visualizations generated using IGV^65^. The code is available at https://github.com/broadinstitute/Klebs_meroR_gwas.git or from the authors upon request.

## Acknowledgements

We thank Thuy-Lan Lite and Tatum Mortimer for their valuable advice during this project. We thank Sahil Sood for assistance in candidate selection. Research reported in this publication was supported by the National Institute Of Allergy And Infectious Diseases of the National Institutes of Health under Award Number R21AI199686. The content is solely the responsibility of the authors and does not necessarily represent the official views of the National Institutes of Health.

