## Supplementary Information for "Genome-Wide Association Study Identifies Novel Genetic Variants Linked to Meropenem Resistance in *Klebsiella pneumoniae*"

For the first test, we conditioned on the presence of the 22 resistance-associated unitigs that mapped to the *hsdM-hsdS-hsdR-IsKpn25* genes in addition to the presence of carbapenemase genes by adding them as additional covariates to the LMM, independently testing each of the 20 resistance-associated unitigs that mapped to the *umuC* gene once more. We observed that 19 of the 20 unitigs we tested again were still significant with positive effect sizes (Supplementary Fig 2a) strengthening the case for the resistance-associated *umuC* allele causing increased resistance to meropenem by an undetermined mechanism.

For the second test, we conditioned on the presence of the 20 resistance-associated unitigs that mapped to *UmuC* gene in addition to the presence of carbapenemase genes by adding them as additional covariates to the linear mixed model, independently testing each of the 22 resistance-associated unitigs that mapped to the *hsdM-hsdS-hsdR-IsKpn25* genes once more. We observed that none of the 22 unitigs we tested were statistically significant (Supplementary Fig 2b). However, when we tested a single variable representing the presence and absence of the entire *hsdM-hsdS-hsdR-IsKpn25* element by grouping together all 22 resistance-associated unitigs, while conditioning on the 20 resistance-associated *UmuC* unitigs and the carbapenemase genes using a LMM as before (see Methods), the single variable representing the entire element was statistically significant ($\beta = 0.932$, *p*-value = 0.017). We surmised this was because testing the entire *hsdM-hsdS-hsdR-IsKpn25* element by grouping together individual unitigs increased the power of the conditional LMM compared to testing each individually by implicitly accounting for LD between the *hsdM-hsdS-hsdR-IsKpn25* resistance-associated unitigs.

To determine whether there was a synergistic effect between *hsdM-hsdS-hsdR-IsKpn25* and *umuC* we conducted another statistical test, this time conditioning on the presence of carbapenemases, *umuC* and *hsdM-hsdS-hsdR-IsKpn25* by adding them as additional covariates, testing an interaction variable for *umuC* and *hsdM-hsdS-hsdR-IsKpn25* (see Methods) using a linear mixed model. The interaction variable was statistically significant with a negative effect size ($\beta = -1.16$, *p*-value = 0.007). We conjectured that this implied the individual effects from *UmuC* and *hsdM-hsdS-hsdR-IsKpn25* are not linearly additive. This could be explained by redundance in the biological mechanisms by which each confers resistance or the saturation of the assay used to measure resistance itself.
